# An Explainable and Robust Deep Learning Approach for Automated Electroencephalography-based Schizophrenia Diagnosis

**DOI:** 10.1101/2023.05.27.542592

**Authors:** Abhinav Sattiraju, Charles A. Ellis, Robyn L. Miller, Vince D. Calhoun

## Abstract

Schizophrenia (SZ) is a neuropsychiatric disorder that affects millions globally. Current diagnosis of SZ is symptom-based, which poses difficulty due to the variability of symptoms across patients. To this end, many recent studies have developed deep learning methods for automated diagnosis of SZ, especially using raw EEG, which provides high temporal precision. For such methods to be productionized, they must be both explainable and robust. Explainable models are essential to identify biomarkers of SZ, and robust models are critical to learn generalizable patterns, especially amidst changes in the implementation environment. One common example is channel loss during EEG recording, which could be detrimental to classifier performance. In this study, we developed a novel channel dropout (CD) approach to increase the robustness of explainable deep learning models trained on EEG data for SZ diagnosis to channel loss. We developed a baseline convolutional neural network (CNN) architecture and implement our approach as a CD layer added to the baseline (CNN-CD). We then applied two explainability approaches to both models for insight into learned spatial and spectral features and show that the application of CD decreases model sensitivity to channel loss. The CNN and CNN-CD achieved accuracies of 81.9% and 80.9% on testing data, respectively. Furthermore, our models heavily prioritized the parietal electrodes and the α-band, which is supported by existing literature. It is our hope that this study motivates the further development of explainable and robust models and bridges the transition from research to application in a clinical decision support role.

## I. Introduction

In recent years, the use of machine and deep learning methods in studies for automated diagnosis of neuropsychiatric disorders has increased, illustrating the potential for widespread use of artificial intelligence (AI) in clinical diagnostic efforts in the future [1]. This could be particularly useful for disorders like schizophrenia (SZ), in which diagnosis is based on symptoms that often vary across individuals and overlap with related disorders [2]. Many such studies have used electroencephalography (EEG) [3], [4], which provides insight into brain activity dynamics with high temporal precision. For machine and deep learning methods to transition from research to clinical implementations, methods for ensuring model robustness and explainability are needed to guarantee that learned patterns are both identifiable and generalizable. In this study, we implement a novel approach for improved robustness to the common issue of EEG channel loss on an explainable deep learning classifier.

Many studies have developed models for automated EEG-based diagnosis of neuropsychiatric disorders like major depressive disorder [5], [6], Alzheimer’s disease [7], SZ [3], [4], and more. SZ, in particular, has symptoms that overlap with other disorders and is challenging to diagnose [2]. As such, it presents a need for the development of clinical decision support systems that identify key generalizable biomarkers. Earlier EEG-based studies for automated SZ diagnosis used manual feature engineering with machine learning models [8]. Feature engineering is challenging and time-consuming, and recent EEG-based studies have made the transition to deep learning methods capable of automated feature extraction. For such methods to be applicable in clinical settings, they must be both explainable and robust.

Explainable models are important from both scientific and clinical perspectives. They have the potential to identify biomarkers that contribute to a better understanding of disorders and that clinicians could eventually use in practice [9]. Furthermore, clinicians have an ethical responsibility to explain recommendations of clinical decision support systems to patients [10]. There are multiple types of explainability methods that have been recently developed for raw EEG classifiers. (1) Spatial explainability methods indicate the relative importance of different EEG channels to model performance [11], [12], and (2) spectral explainability methods indicate the relative importance of different frequency bands to model performance [13], [14]. The results of these explainability methods can verify patterns identified in literature but also find novel features as well. Spatial explainability methods are especially useful in analyzing model robustness to channel perturbation.

Robust models are those that have learned generalizable patterns. Innovations such as dropout and spatial dropout reduce model overfitting and improve generalizability to multiple datasets, allowing classifiers to learn the underlying associations between features and responses. These approaches have been used in deep-learning-based classification of EEG data [3], [15]. Robust models can also handle changes in the implementation environment. One common change in the environment of EEG-based classifiers is channel loss, which is associated with electrodes that are broken or improperly attached. A model that is not robust to channel loss may see a decrease in performance if important channels are lost, which motivates the need for methods that reduce model sensitivity to channel loss and increase overall robustness [16].

In this study, we apply a novel channel dropout (CD) approach to a convolutional neural network (CNN) trained on EEG to increase model robustness to channel loss. We then apply two explainability approaches to identify key features learned by the models and illustrate the effectiveness of CD. We also find that our models learned features associated with SZ that have been identified in existing literature. To the best of our knowledge, our study represents the first study with a combined focus on explainability and robustness of deep learning models trained on raw EEG for SZ diagnosis and a step towards the implementation of deep learning in a clinical decision support role for SZ diagnosis.

## II. Methods

### A. Data Preprocessing

We utilized a scalp EEG dataset presented in [17]. The dataset contains 14 individuals with SZ and 14 healthy controls (HCs), collected at the Institute of Psychiatry and Neurology in Warsaw, Poland. All participants gave written, informed consent. Recordings utilized 64 electrodes placed in the standard International 10-20 system and were performed during a 15-minute resting state at 250 Hz. We only used 19 electrodes (Fp1, Fp2, F7, F3, Fz, F4, F8, T3, C3, Cz, C4, T4, T5, P3, Pz, P4, P6, T6, O1, and O2) like other studies [4]. We used a moving window approach with a 2.5-second step size to separate the recordings into 25-second epochs. We then upsampled the HCs to balance the data, detrended all samples by fitting to a polynomial basis of degree 5 [18], and channel-wise z-scored all samples. Our final dataset consisted of 6,036 SZ and 6,036 HC samples.

### B. Model Development

We developed a baseline CNN using an architecture that was adapted from a prior study which utilized a similar EEG data format, making the performant architecture easily adaptable [4]. To emphasize generalizability, we included MaxNorm regularization and batch normalization [3]. We trained the baseline CNN and the CNN with an added CD layer (CNN-CD) as depicted in Fig. 1. The CD layer zeroed out channels at a rate of 15% across all training samples.

**Fig 1.**
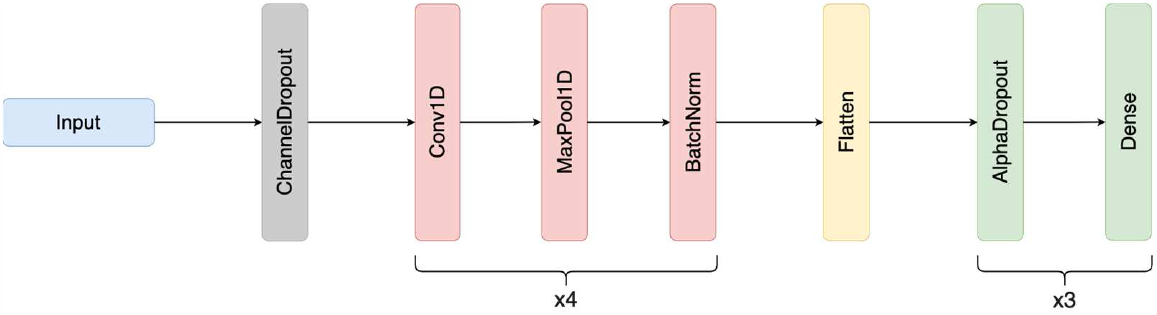
CNN-CD Architecture. The ChannelDropout layer has a dropout rate of 0.15. The convolution block has 4 1D Convolutional layers (Conv1D). All Conv1D layers have a stride of 1. The first Conv1D layer has 5 filters of length 10. The second Conv1D layer has 10 filters of length 15. The third Conv1D layer has 15 filters of length 15. The fourth Conv1D layer has 15 filters of length 10. Each Conv1D layer is followed by a Max Pooling layer with a pool size of 2 and stride of 2 and a Batch Normalization layer. The classifier portion of the CNN contains 3 Dense layers with 64 nodes, 32 nodes, and 1 node. All Dense layers are preceded by an AlphaDropout layer with a dropout rate of 0.5. All Conv1D and Dense layers have Exponential Linear Unit (ELU) activations, He Normal weight initialization, and MaxNorm regularization with a maximum value of 1. The final Dense layer has a Sigmoid activation and Glorot Normal weight initialization. The CNN architecture is the same as the CNN-CD without the ChannelDropout layer.

In our training process, we used a 10-fold cross-validation strategy where approximately 72%, 14%, and 14% of the data were assigned to the train, validation, and test sets, respectively. Data was split by subject, rather than sample, ensuring that the model evaluated subjects in the test set that were unseen in the training set. This enables test performance to more reliably represent the generalizability of the model. Like in [3], we applied data augmentation to the training set for each fold prior to training to ensure the models learned features important for SZ classification amidst noise. We generated 3 copies of the training data. We applied gaussian noise with a mean (µ) of 0 and a standard deviation (σ) of 0.7 to one copy and sinusoidal noise with a frequency of 2 Hz and amplitude of 0.3 to the other copy. We left the final copy unchanged and used all copies in our augmented training set.

Both models were trained for 50 epochs per fold with a batch size of 256, binary cross-entropy loss, and an Adam optimizer with a learning rate of 0.001. The loss was weighted to account for any class imbalances caused by the train-validation-test split. Across all epochs, we tracked accuracy (ACC), sensitivity (SENS), specificity (SPEC), and balanced accuracy (BACC) to measure performance. For each fold, we used the model from the epoch with the maximal validation BACC to evaluate the test set performance. We calculated the µ and σ of all metrics across folds.

### C. Explainability Approaches

We applied two explainability approaches in this study. In our first approach, we ablated individual EEG channels in an approach mimicking the line-related noise found in EEG recordings that was first used in multimodal electrophysiology [11]. In our second approach, we ablated individual frequency bands within each channel. This approach resembled the EasyPEASI spectral explainability approach [13].

#### 1) Spatial Explainability

For insight into the importance of spatial features learned by both models, we applied an ablation approach along the channel axis with the following steps: (1) We calculated the model BACC on the test data. (2) We iterated through all 19 channels and ablated each channel individually with sinusoidal noise with a frequency of 60 Hz and amplitude of 0.1. (3) We calculated the model BACC on the perturbed test data after each iteration and measured the percent change.

#### 2) Spatial-Spectral Explainability

For insight into the importance of spatial and spectral features learned by both models, we applied a perturbation approach with the following steps: (1) We calculated the model BACC on the test data. (2) We performed a Fast Fourier Transform (FFT) to convert each sample to the frequency domain. (3) For each of the 19 channels we iterated through, we substituted the spectral coefficients within each frequency band individually with Gaussian noise with the same μ and σ as the substituted spectral coefficients. We iterated through the following frequency bands in Hertz (Hz) for each channel: δ (0 – 4 Hz), θ (4-8 Hz), α (8 – 12 Hz), β (12 – 25 Hz), γ_lower_ (25 – 55 Hz), and γ_upper_ (65 – 125 Hz). (4) We converted the perturbed samples back to the time domain with an inverse FFT, and (5) we calculated the percent change in model BACC after each perturbation. This approach allowed us to understand the importance of various frequency bands within each channel to model performance.

## III. Results and Discussion

In this section, we discuss the performance and explainability results of both models.

### A. Model Performance

Table 1 shows the ACC, SENS, SPEC, and BACC μ and σ of our CNN and CNN-CD. Both architectures performed above 75% across all metrics. The mean CNN performance was slightly greater than that of the CNN-CD across all metrics except SPEC. However, the CNN had a greater variability in performance than the CNN-CD across all metrics except SPEC. Both models performed at or better than the CNN in [3], which was trained with the same subject-based cross-validation approach. The SENS and SPEC of our models were lower than that of [4]. However, that study used a cross-validation approach that allowed samples from the same subjects to be distributed across training, validation, and test sets simultaneously. As addressed in [19], that cross-validation approach can inflate model performance. Our cross-validation approach allowed us to gain a greater understanding of the generalizability of model results.

**TABLE 1.**
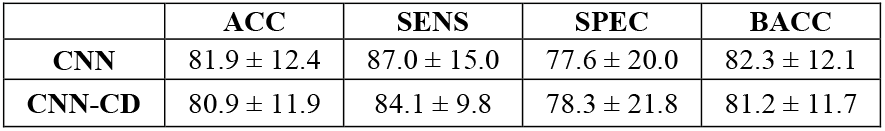
Classification Performance Results

### B. Spatial Explainability

Fig. 2 shows the effects of ablating individual EEG channels on the CNN and CNN-CD test performance. Perturbation of the parietal and central electrodes (i.e., PZ, P7, C4, CZ) are associated with larger changes in BACC of the CNN relative to other electrodes as seen in Fig. 2a, indicating their importance to the CNN in SZ classification. The importance of these channels is supported by existing literature. The identified electrodes are located near the parietal lobe, which has reduced structural connectivity in SZs [20]. On the other hand, perturbation of parietal and central electrodes, especially PZ, are associated with smaller changes in BACC of the CNN-CD relative to the CNN as seen in Fig. 2b. This lends credibility to CD causing the model to distribute importance more evenly across channels rather than relying on a subset of channels. This is important for maintaining model performance in a clinical setting where key channels may be lost during recording.

**Fig 2.**
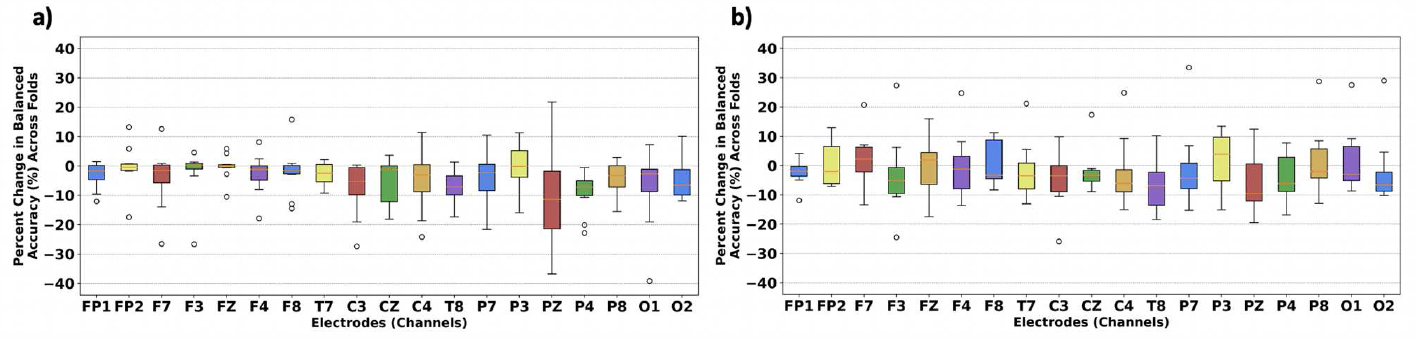
Plots of Spatial Importance. Panels a) and b) show the importance of each channel to the CNN and CNN-CD, respectively. The percent change in BACC after perturbation across folds is shown on the y-axis, and the x-axis indicates specific channels that were perturbed.

### C. Spatial-Spectral Explainability

Fig. 3 illustrates the effects of perturbing frequency bands within channels on test performance for both models. For the CNN, δ and α were most important, especially in association with central electrodes (i.e., C3, C4) and parietal electrodes (i.e., PZ, P3, P4) respectively. The importance of the identified parietal electrodes to the CNN fits with our spatial explainability results in Figure 2a. Additionally, these results are supported by past literature, as C3 has been found to have less complexity in SZs than in HCs [21]. Slower α oscillations have also been associated with cognitive deficits in SZs [22], and δ has been found to have increased oscillations in SZs [23]. Similarly, the CNN-CD found α to be most important in association with PZ and P4. Note that unlike the CNN, the CNN-CD prioritized F3 and F7 in association with δ. Those electrodes are located near the frontal lobe, and SZ is known to impair executive function, which is associated with the prefrontal cortex [24]. Due to CD, the distribution of importance for the CNN-CD is less concentrated than that of the CNN. Our spatial explainability approach shows the high relative importance of PZ and F3 to the CNN-CD in Figure 2b, illustrating how both explainability approaches align in relative channel importance to the models.

**Fig 3.**
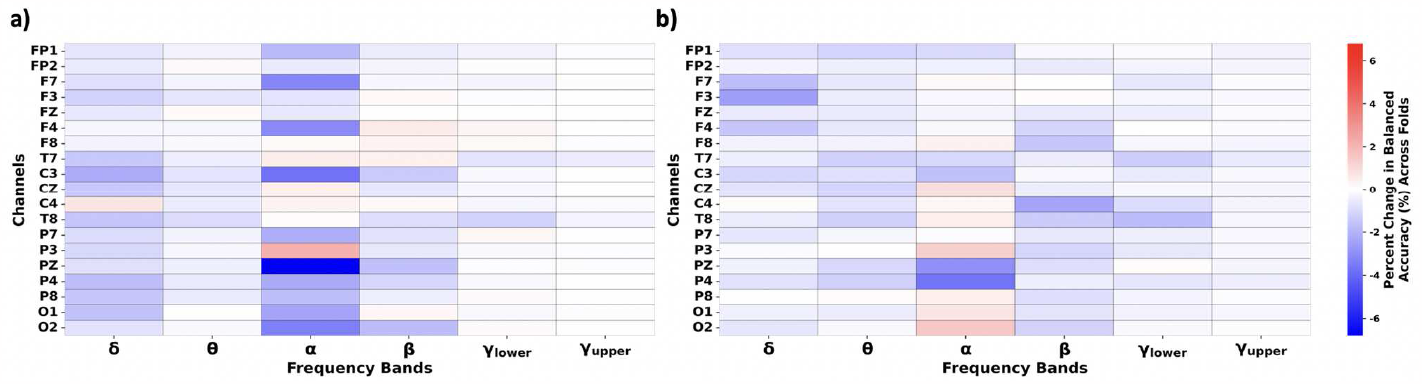
Plots of Spatial-Spectral Importance. Panels a) and b) show the importance of each channel-frequency-band combination to the CNN and CNN-CD, respectively. The percent change in balanced accuracy after perturbation across folds is reflected on the heatmap, and the color bar to the right of panel b) is shared by both panels.

### D. Limitations and Next Steps

In this study, we focused on the effect of CD on a CNN architecture. It would be useful to examine the effect of CD on other CNN or CNN-LSTM architectures that have been used in past studies [3], [4]. It would be beneficial to explore the effects of multiple CD rates on model performance and robustness to channel perturbation. We could also examine the effect of the CNN-CD on other EEG SZ datasets to further assess the generalizability of learned patterns. It would also be interesting to examine the effect of CD on models trained on EEG data for detection of major depressive disorder, Alzheimer’s disease, and other prominent disorders to which past studies have applied deep learning methods for automated diagnosis [6], [7].

## IV. Conclusion

In this study, we demonstrated the utility of a novel CD approach in reducing sensitivity to EEG channel loss for an explainable CNN architecture for SZ diagnosis. We applied two explainability approaches for insight into spatial and spectral features learned by the CNN and CNN-CD. We found that both models heavily prioritized parietal electrodes and the α-band. Importantly, the CNN-CD was less sensitive to spatial perturbation due to CD forcing the model to learn from more channels rather than overfitting on a specific subset of channels. Our explainability findings are consistent with previous studies [20]–[24], and it is our hope that this study will help further the development of robust, explainable models that bridge the gap between research and production.

